# Sphingosine synergizes with polymyxin antibiotics to kill Gram-negative bacteria

**DOI:** 10.1101/2025.07.28.667204

**Authors:** Jacob R. Mackinder, Meghan Quinlan, Matthew J. Wargo

## Abstract

Antimicrobial resistance is an increasing threat to global health. However, there is a limited set of antibiotics that are effective against drug-resistant Gram-negative bacteria like *Pseudomonas aeruginosa*. One strategy to enhance the efficacy and longevity of existing antibiotics is by combining them with non-traditional antimicrobial adjuvants. Here, we examined if the host-derived antimicrobial lipid sphingosine could enhance the efficacy of a panel of antibiotics against *P. aeruginosa in vitro*. We found that sphingosine displayed strong synergy with the polymyxin antibiotics, polymyxin B and colistin, to inhibit growth of and kill *P. aeruginosa,* but did not significantly alter the efficacy of other tested antibiotic classes. The addition of sphingosine reduced the MIC of polymyxin B and colistin from 0.5 µg/mL to 0.031 µg/mL and 8 µg/mL to 0.5 µg/mL, respectively. This combination of sphingosine and polymyxin B synergized to inhibit the growth and survival of *Klebsiella pneumoniae* as well. In addition to sphingosine, we found that the sphingoid bases sphinganine (dihydrosphingosine) and phytosphingosine also enhanced the activity of polymyxins. Overall, these findings demonstrate that sphingosine is a potent adjuvant for polymyxins, and that the sphingosine- polymyxin combination is capable of killing *P. aeruginosa* and *K. pneumoniae* while using relatively low concentrations of polymyxin. This study may help in the development of new antimicrobial therapies for the treatment of Gram-negative bacterial infections.

**Importance:** Antibiotic resistance is an increasing threat to global health and there is a dire need to develop new therapies to treat multidrug-resistant infections. Here we show that sphingosine, a eukaryotic-derived antimicrobial lipid, synergizes with polymyxin antibiotics to inhibit the growth and survival of the Gram-negative bacteria, *P. aeruginosa* and *K. pneumoniae*. Thus, sphingosine has potential as an antimicrobial adjuvant for combinatorial therapies with polymyxins to treat Gram-negative infections.

## Introduction

Antibiotic resistance is a major problem with the number of antibiotic-resistant infections increasing [1]. Many antibiotic-resistant infections can be attributed to a specific group of bacteria referred to as the “ESKAPE” pathogens, which include *Enterococcus faecium*, *Staphylococcus aureus*, *Klebsiella pneumoniae*, *Acinetobacter baumannii*, *Pseudomonas aeruginosa* and other members of the Enterobacterales group [2, 3]. ESKAPE pathogens have been designated “priority status” by the World Health Organization (WHO) with the goal to increase research, discovery, and development of new antibiotics to combat these pathogens.

*P. aeruginosa* is a Gram-negative opportunistic bacterial pathogen capable of infecting numerous body sites including the lungs, bloodstream, urinary tract, burns, and surgical wounds [4–7]. Its success as an opportunist is partially due to its ability to rapidly adapt to environmental cues and stressors, including antibiotic exposure [8–11]. *P. aeruginosa* has a variety of intrinsic and acquired antimicrobial resistance mechanisms which include biofilm formation, reduced outer membrane permeability, efflux systems, and drug-inactivating enzymes [10–14]. Multi-drug resistant (MDR) strains of *P. aeruginosa* are often associated with hospital- and ventilator-acquired pneumonia, bacteremia, and chronic lung infections in individuals with cystic fibrosis (CF) and chronic obstructive pulmonary disease [4–6, 10, 15–17]. From 2015-2017, the National Healthcare Safety Network found that 26% of *P. aeruginosa* isolates from ICU patients were resistant to carbapenems, 26.5% were resistant to extended-spectrum cephalosporins, 27.1% were resistant to fluoroquinolones, and 18.6% were resistant to three or more antibiotics [17]. Although most *P. aeruginosa* infections can be treated with currently available antibiotics, the continued increase of antibiotic resistance rates combined with limited development of new antimicrobials is a dire situation [18]. Therefore, there is an urgent need to develop novel antimicrobial strategies, one of which is through the combination of existing antibiotics with non-traditional adjuvants.

Antimicrobial combinations that repurpose existing antibiotics offer a sustainable and cost-effective alternative to the discovery and development of novel antibiotics. Combinatorial therapies can result in improved outcomes over monotherapy when the primary drug alone has a high likelihood of inducing resistance (*e.g*. fosfomycin), the pathogen has a high rate of developing resistance (*e.g. P. aeruginosa*), or when there are limited treatments for highly resistant infections in vulnerable patients [10, 11, 19, 20]. The best studied combinatorial therapies for drug resistant infections are β-lactam antibiotics paired with β-lactamase inhibitors; however, host-derived antimicrobial factors have also been explored [21–25]. Sphingolipids are a class of lipids primarily found in eukaryotes that have a variety of biological functions, including immune signaling and antimicrobial activity [25–27]. The sphingoid base sphingosine, the core moiety of most sphingolipids, has potent antimicrobial properties, capable of inhibiting growth and killing both Gram-negative and Gram-positive bacteria [26, 28–30]. In the case of Gram-negative bacteria, sphingosine destabilizes the outer membrane by binding to cardiolipin, resulting in increased cardiolipin turnover and degradation with eventual leakage of ATP and loss of metabolic activity [31, 32]. The anti-infectious role of sphingosine in epithelial tissues have been studied primarily in the contexts of atopic dermatitis and CF, where the reduction of local sphingosine concentrations in the stratum corneum and airway lumen, respectively, leaves these tissues more susceptible to infection [33–36]. Restoring the sphingolipid homeostasis in CF mice through inhalation of aerosolized sphingosine or recombinant acid ceramidase, which releases sphingosine from accumulated ceramide, reduces infection and improves survival outcomes [35–37]. The anti-infectious role of sphingosine led us to test whether this host- compound would be able to synergize with currently available antibiotics to inhibit growth of *P. aeruginosa*.

## RESULTS

### Screen of antibiotics for altered activity with sphingosine

To test if sphingosine could be used as an antimicrobial adjuvant, we measured *P. aeruginosa* growth in a panel of CF-relevant antibiotics, covering five antibiotic classes: aminoglycosides (gentamicin and tobramycin), β-lactams (aztreonam and ceftazidime), cyclic polypeptides (polymyxin-B and colistin), a fluoroquinolone (ciprofloxacin), and a glycopeptide (vancomycin) (**Table 1**). *P. aeruginosa* cultures were grown in minimal media containing serial dilutions (1:1, v:v) of antibiotic with or without 20, 70, or 200 µM sphingosine. These sphingosine concentrations were selected because they do not inhibit wildtype *P. aeruginosa* growth [38] and are found in physiologically-relevant environments of *P. aeruginosa* infection: airway surface liquid (20 µM), nasal mucosa (70 µM), and the stratum corneum of the skin (200 µM) [33, 39–45].

**Table 1.** Mean MICs of antibiotics screened alone or with 200 µM sphingosine

| Class | Antibiotic | Target | PA14<br>MIC<br>(µg/mL) | PA14<br>MIC<sub>combination</sub><br>(µg/mL) | PAO1<sup>a</sup><br>MIC<br>(µg/mL) | PAO1<sup>a</sup><br>MIC<sub>combination</sub><br>(µg/mL) |
| --- | --- | --- | --- | --- | --- | --- |
| Aminoglycoside | Gentamicin | Ribosome | 2 | 2 | 2 | 4 |
|  | Tobramycin | Ribosome | 1 | 2 | 1 | 2 |
| β-lactam | Aztreonam | Peptidoglycan | 1 | 1 | - | - |
|  | Ceftazidime | Peptidoglycan | 4 | 4 | - | - |
| Fluoroquinolone | Ciprofloxacin | Topoisomerase | 1 | 2 | - | - |
| Glycopeptide | Vancomycin | Peptidoglycan | >512 | >512 | - | - |
| Polymyxins | Polymyxin B | Outer membrane | 0.5 | 0.03125 | 0.5 | 0.125 |
|  | Colistin | Outer membrane | 8 | 0.5 | 8 | 2 |

<sup>a</sup>, PAO1 was not tested against β-lactam, fluoroquinolone, or glycopeptide antibiotics.
-, not calculated.

Regardless of the concentration, sphingosine did not alter the MICs of aztreonam, ceftazidime, or vancomycin (**Table 1, Supplemental Figures 1-5**). The presence of sphingosine (20, 70 and 200 µM) had a small effect on the MIC of aminoglycosides, shifting it from 1 µg/mL to 2 µg/mL or 4 µg/mL for tobramycin and gentamicin, respectively (**Supplemental Figures 1-2**). Likewise, 200 µM of sphingosine moved the MIC of ciprofloxacin from 1 µg/mL to 2 µg/mL (**Supplemental Figure 3**). Conversely, the addition of sphingosine to the two polymyxin antibiotics, polymyxin B and colistin (polymyxin E), substantially reduced the MIC of both drugs. The MIC of polymyxin-B dropped from 0.5 µg/mL alone, to 0.125 µg/mL and 0.031 µg/mL with the addition of 20 µM and either 70 µM or 200 µM sphingosine, respectively (**Figure 1A**, **Table 1**), thus either 70 µM and 200 µM sphingosine reduced the polymyxin-B MIC 16-fold. The addition of 20 µM sphingosine reduced the MIC of colistin from 8 µg/mL to 2 µg/mL, while the addition of either 70 µM and 200 µM sphingosine reduced the MIC of colistin to 0.5 µg/mL (**Figure 1B**, **Table 1**), the latter representing approximately a 16-fold reduction in colistin concentration. In addition to PA14, 200 µM sphingosine enhanced polymyxin antibiotics against PAO1 with a shift in the MIC of polymyxin B from 0.5 µg/mL to 0.125 µg/mL and the MIC of colistin dropped from 8 µg/mL to 2 µg/mL (**Figure 1C-D**, **Table 1)**. Because PA14 displayed the strongest level of synergy it was used for subsequent experiments.

**Figure 1.**
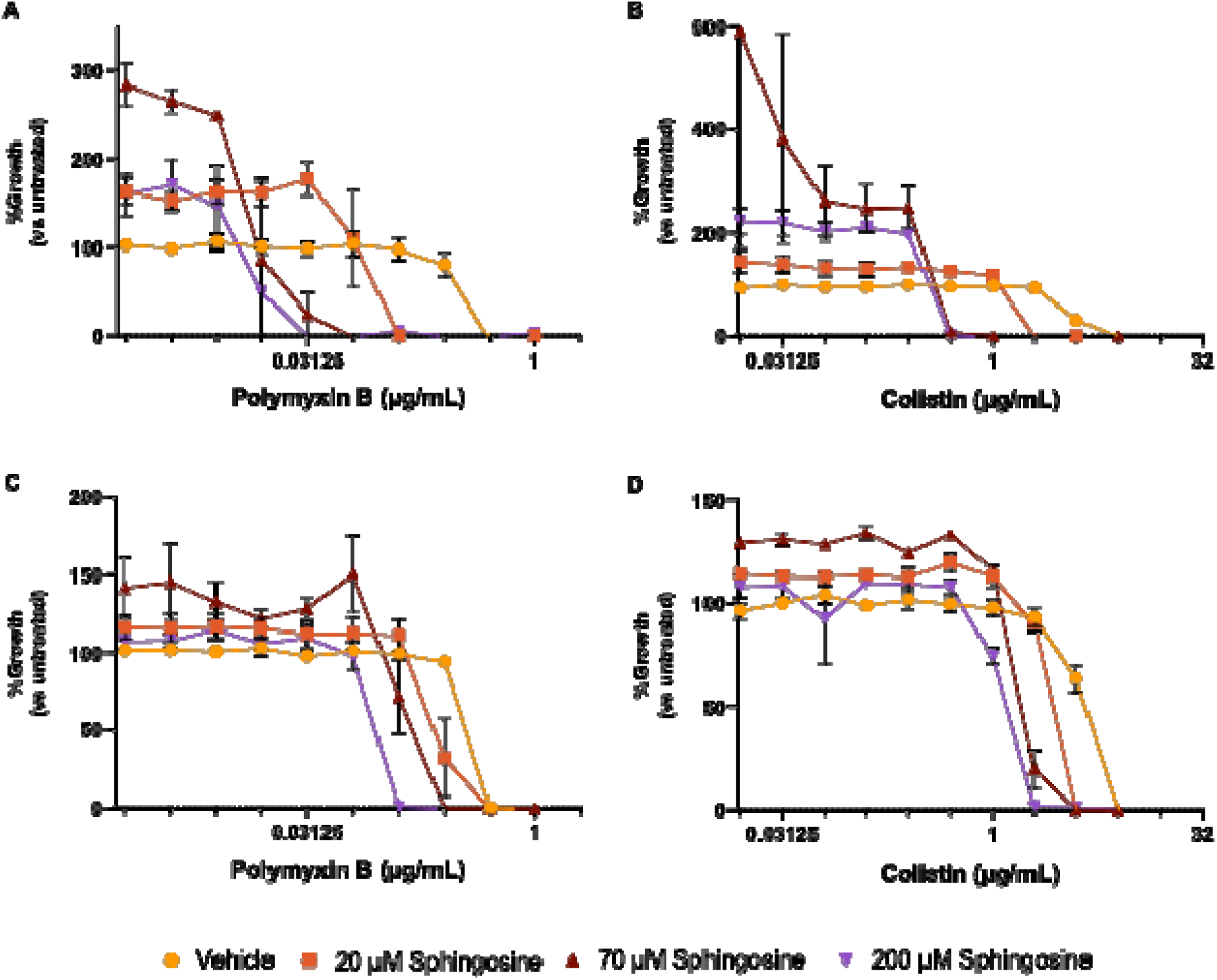
Sphingosine reduces MIC of polymyxin antibiotics. Dose-effect curves of PA14 in increasing concentrations of polymyxin B **(A)** or colistin **(B)** alone or with 20 µM, 70 µM, and 200 µM sphingosine. Dose-effect curves of PAO1 in increasing concentrations of polymyxin B **(C)** or colistin **(D)** alone or with 20 µM, 70 µM, and 200 µM sphingosine. Data points denote means summarizing three independent experiments and error bars represent the standard error of mean.

**Figure 2.**
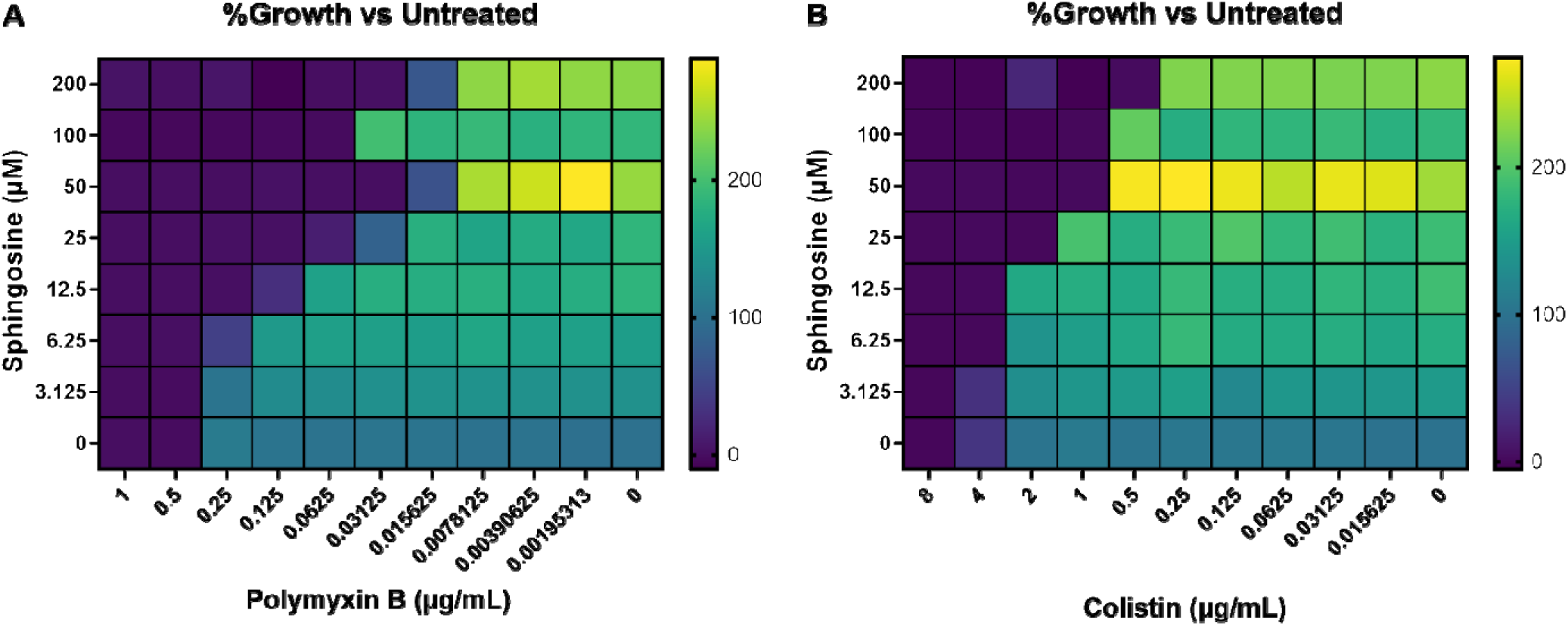
Checkerboard assays of *P. aeruginosa* growth with combinations of sphingosine and polymyxin antibiotics. Heatmaps representing percent PA14 growth in polymyxin B **(A)** and colistin **(B)** with a series of sphingosine concentrations. Presented as the mean percent-growth relative to the untreated control from three independent experiments.

### Sphingosine and polymyxin antibiotics synergize to inhibit *P. aeruginosa* growth

To determine if the enhanced antibiotic activity of polymyxins and sphingosine represented antibiotic synergy, we conducted checkerboard assays with serial dilutions of both compounds. Checkerboard assays were used to calculate the fractional inhibitory concentration index (FICI) of the combinations (**Equation 1**). For categorization of FIC values, we used the definitions of ≤0.5 as synergistic, 0.5-1.0 as additive, 1-2 as unrelated, and >2 as antagonistic [46]. Sphingosine enhanced efficacy of both polymyxin antibiotics at multiple concentrations.

Synergy between sphingosine and polymyxin B was observed when 50 µM sphingosine was combined with 0.031 µg/mL polymyxin B and when 25 µM sphingosine was used with 0.125 µg/mL polymyxin B resulting in FICIs of 0.26, and 0.35, respectively (**Figure 2A**, **Table 2**). Colistin combined with sphingosine resulted in a FICI ranging from 0.325-0.35 when using 50 µM sphingosine and 1 µg/mL colistin, or 25 µM sphingosine with 2 µg/mL colistin, respectively (**Figure 2B**, **Table 2**).

**Table 2.** MICs and FICIs of sphingoid bases and polymyxin antibiotics.

| Strain | Sphingoid Base |  |  | Antibiotic |  |  | FICI Combination<sup>a</sup> | Interpretation<sup>b</sup> |
| --- | --- | --- | --- | --- | --- | --- | --- | --- |
|  | Compound | MIC (μM) | MIC<sub>combination</sub> (μM) | Compound | MIC (μg/mL) | MIC<sub>combination</sub> (μg/mL) |  |  |
| *P. aeruginosa* PA14 | Sphingosine | >250 | 200 | Polymyxin B | 0.5 | 0.03125 | 0.8625 | Additive |
|  |  |  | 100 |  |  | 0.0625 | 0.525 | Additive |
|  |  |  | 50 |  |  | 0.03125 | 0.2625 | Synergistic |
|  |  |  | 25 |  |  | 0.125 | 0.35 | Synergistic |
|  |  |  | 12.5 |  |  | 0.25 | 0.55 | Additive |
|  |  |  | 200 | Colistin | 8 | 0.5 | 0.8625 | Additive |
|  |  |  | 100 |  |  | 1 | 0.525 | Additive |
|  |  |  | 50 |  |  | 1 | 0.33 | Synergistic |
|  |  |  | 25 |  |  | 2 | 0.35 | Synergistic |
|  |  |  | 12.5 |  |  | 4 | 0.55 | Additive |
|  |  |  | 6.25 |  |  | 4 | 0.525 | Additive |
|  | Sphinganine | >250 | 50 | Polymyxin B | 0.5 | 0.25 | 0.7 | Additive |
|  | Phytosphingosine | 50 | 25 | Polymyxin B | 0.5 | 0.125 | 0.75 | Additive |
|  |  |  | 12.5 |  |  | 0.25 | 0.75 | Additive |
| *K. pneumoniae* MGH78578 | Sphingosine | >250 | 200 | Polymyxin B | 0.5 | 0.0078125 | 0.815625 | Additive |
|  |  |  | 100 |  |  | 0.015625 | 0.43125 | Synergistic |
|  |  |  | 50 |  |  | 0.0078125 | 0.215625 | Synergistic |
|  |  |  | 25 |  |  | 0.03125 | 0.1625 | Synergistic |
|  |  |  | 12.5 |  |  | 0.0625 | 0.175 | Synergistic |
|  |  |  | 6.25 |  |  | 0.25 | 0.525 | Additive |
|  |  |  | 3.125 |  |  | 0.25 | 0.5125 | Additive |
| *K. pneumoniae* KPPR1 | Sphingosine | >250 | 200 | Polymyxin B | 0.5 | 0.015625 | 0.83125 | Additive |
|  |  |  | 100 |  |  | 0.125 | 0.65 | Additive |
|  |  |  | 50 |  |  | 0.0625 | 0.325 | Synergistic |
|  |  |  | 6.25 |  |  | 1 | 2.025 | Antagonistic |
|  |  |  | 3.125 |  |  | 1 | 2.0125 | Antagonistic |

**a**, FICI $\leq 0.5$ , $> 0.5$ and $\leq 1$ , $> 1$ and $\leq 2$ , and $> 2$ , represent synergistic, additive, unrelated, and antagonistic effects, respectively [46].
**b**, Unrelated interactions were not included for brevity.

### Other sphingoid bases enhance the efficacy of polymyxin B against *P. aeruginosa*

In addition to sphingosine, the sphingoid bases sphinganine (dihydrosphingosine) and phytosphingosine display antimicrobial activity [26–30]. Therefore, like sphingosine, we conducted the MIC assays using these two lipids to determine whether they can enhance polymyxin B efficacy. Both sphinganine and phytosphingosine enhanced the antibacterial effects of polymyxin B. The MIC of polymyxin B did not change with 20 µM sphinganine, but in combination with 70 µM and 200 µM the MIC of polymyxin B dropped from 0.5 µg/mL to 0.25 µg/mL (**Figure 3A**). The addition of 20 µM phytosphingosine to polymyxin B reduced the MIC from 0.5 µg/mL to 0.125 µg/mL (**Figure 3B**). *P. aeruginosa* did not grow at any concentration of antibiotic when 70 µM or 200 µM phytosphingosine were present, suggesting that the MIC of phytosphingosine was between 20 and 70 µM.

**Figure 3.**
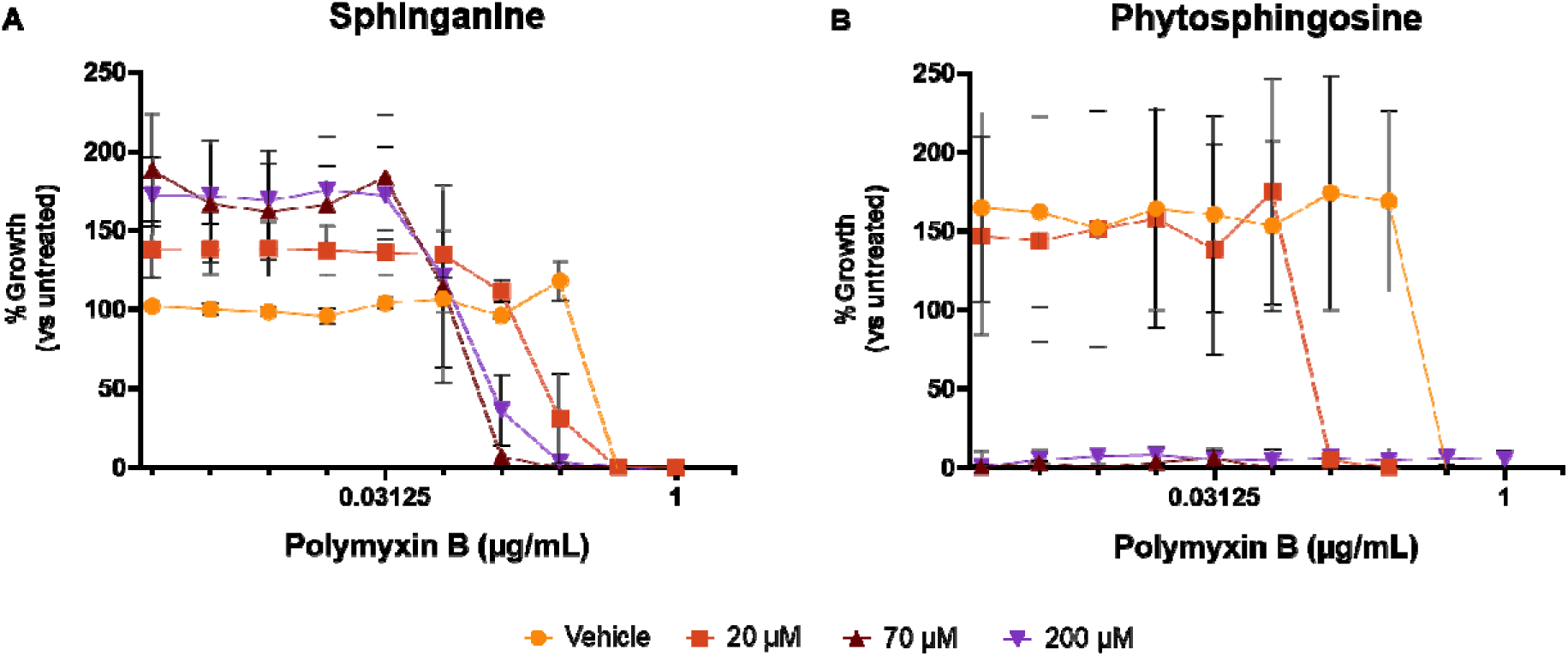
Sphingoid bases reduce polymyxin B MIC. Dose effect curves of PA14 in increasing concentrations of polymyxin B in combination with sphinganine **(A)** and phytosphingosine **(B)**. Data points denote means summarizing three independent experiments and error bars represent the standard error of mean.

Because we observed enhanced antibacterial activity of polymyxin B in combination with sphinganine and phytosphingosine, we conducted checkerboard assays to determine if this enhancement could be defined as synergy with polymyxin B. The combination of 50 µM sphinganine and 0.25 µg/mL polymyxin B had a FICI of 0.70, representing an additive effect (**Figure 4A**, **Table 2**). The checkerboards containing mixtures of polymyxin B and phytosphingosine revealed that 50 µM phytosphingosine alone was sufficient to fully inhibit *P. aeruginosa* growth. The addition of 12.5 µM phytosphingosine to 0.25 µg/mL polymyxin B displayed an additive effect against *P. aeruginosa* growth with a FICI of 0.75, the same FICI was produced from the combination of 25 µM phytosphingosine and 0.125 µg/mL polymyxin B (**Figure 4B**, **Table 2**). FIC values of all sphingoid bases with polymyxins were plotted on <u>isobolograms</u> to illustrate their combined effects on *P. aeruginosa* growth (**Figure 5).**

**Figure 4.**
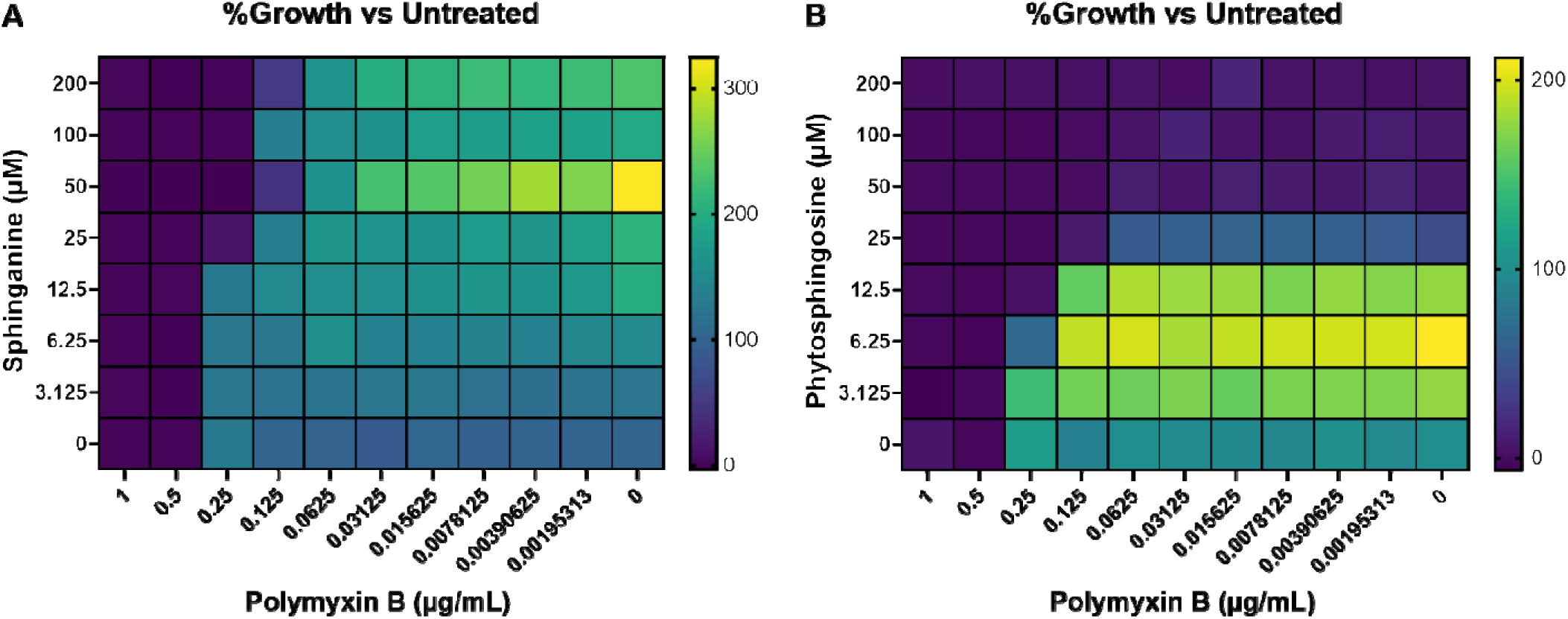
Checkerboard assays of *P. aeruginosa* growth with combinations of sphingoid bases and polymyxin-B. Heatmaps representing percent PA14 growth in different combinations of polymyxin B with sphinganine **(A)** and phytosphingosine **(B)**. Presented as the mean percent-growth relative to the untreated control from three independent experiments.

**Figure 5.**
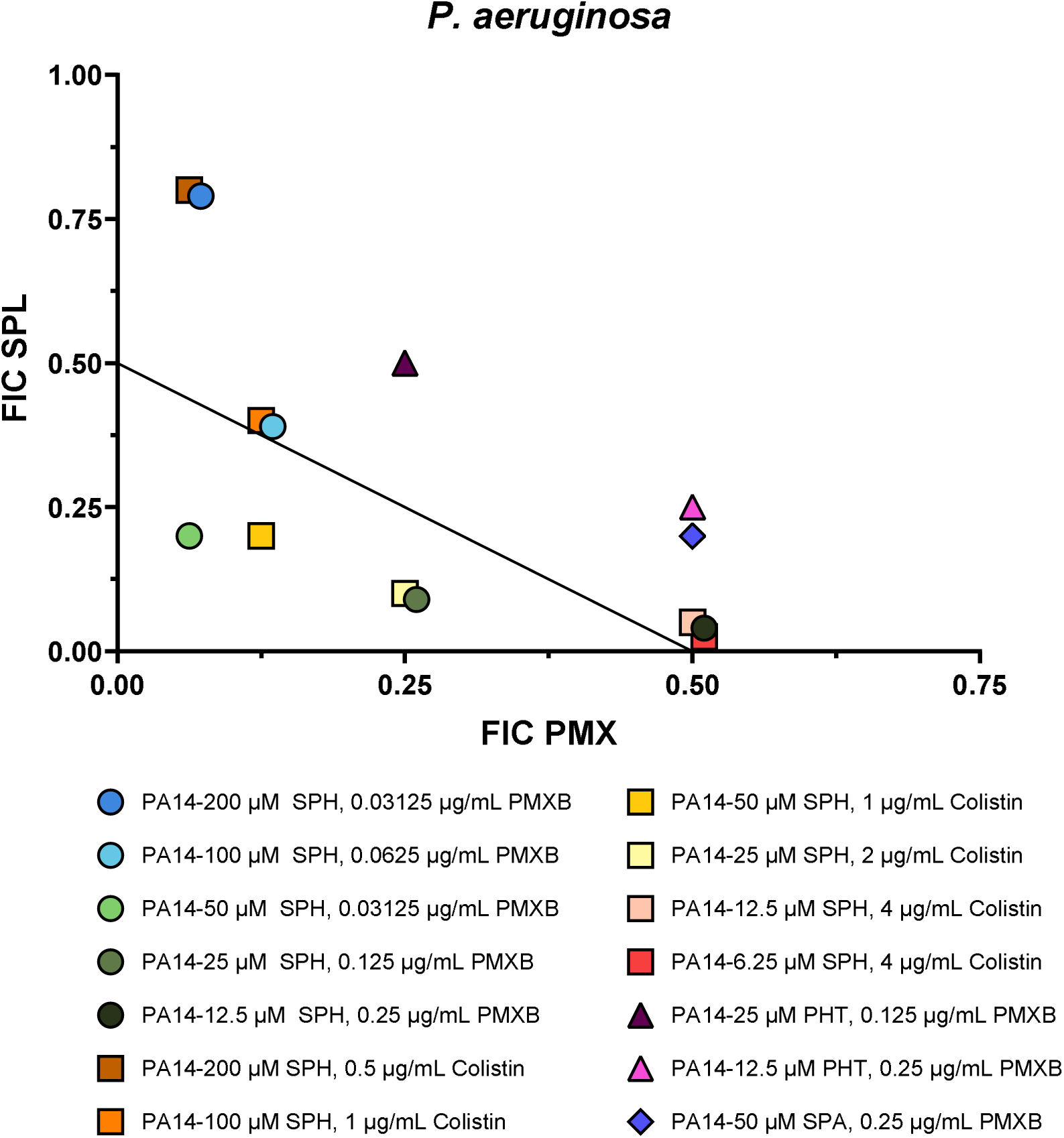
Isobologram of FICI values determined from PA14 checkerboard assays. FIC of sphingoid bases (FIC SPL) were plotted against the FIC of polymyxin antibiotics (FIC PMX). FICI ≤0.5, >0.5 and ≤ 1, >1 and ≤2, and >2, represent synergistic, additive, unrelated, and antagonistic effects, respectively [46].

### Sphingosine synergizes with polymyxin B to inhibit growth of *Klebsiella pneumoniae*

In addition to *P. aeruginosa*, we were interested in the combined activity of sphingosine and polymyxin B on another antibiotic-resistant opportunistic pathogen, *Klebsiella pneumoniae.* Specifically, we focused on combinations of sphingosine and polymyxin B against two *K. pneumoniae* strains: MGH78578 and KPPR1. Both *K. pneumoniae* strains were inhibited by 0.5 µg/mL of polymyxin B alone, but not with sphingosine alone. Growth of MGH78578 was inhibited at multiple combinations of sphingosine and polymyxin B (**Figure 6A**) resulting in a FICI range of 0.16–0.43 (**Figure 7**, **Table 2**). The combination of 25 µM sphingosine and 0.031 µg/mL polymyxin B had the greatest synergistic interaction with an FICI of 0.1625. When KPPR1 was challenged with sphingosine and polymyxin B, 50 µM sphingosine displayed synergy with 0.063 µg/mL polymyxin B with a FICI of 0.325 (**Figures 6B and 7**, **Table 2**).

**Figure 6.**
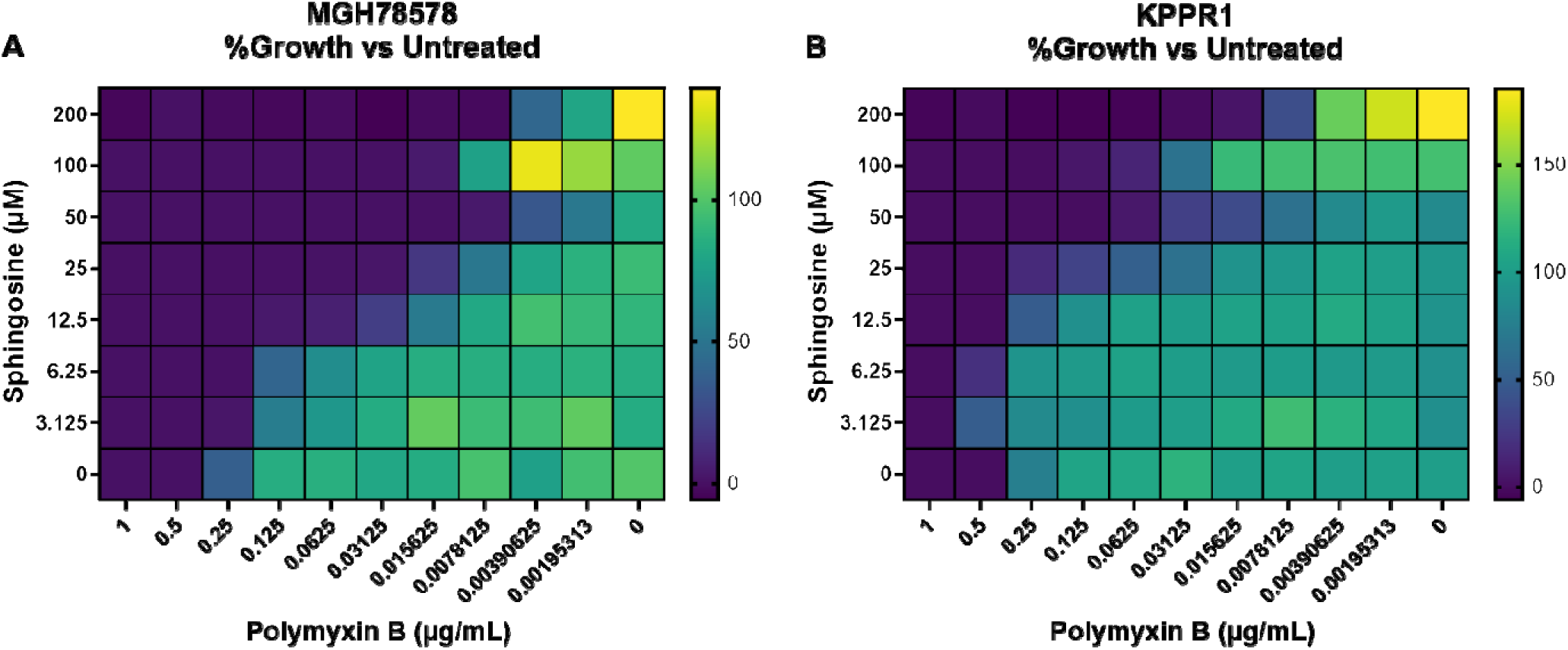
Checkerboard assays of *K. pneumoniae* growth with combinations of sphingosine and polymyxin antibiotics. Heatmaps representing percent growth of *K. pneumoniae* strains MGH78578 **(A)** and KPPR1 **(B)** in different combinations of polymyxin B and sphingosine. Presented as the mean percent-growth relative to the untreated control from three independent experiments.

**Figure 7.**
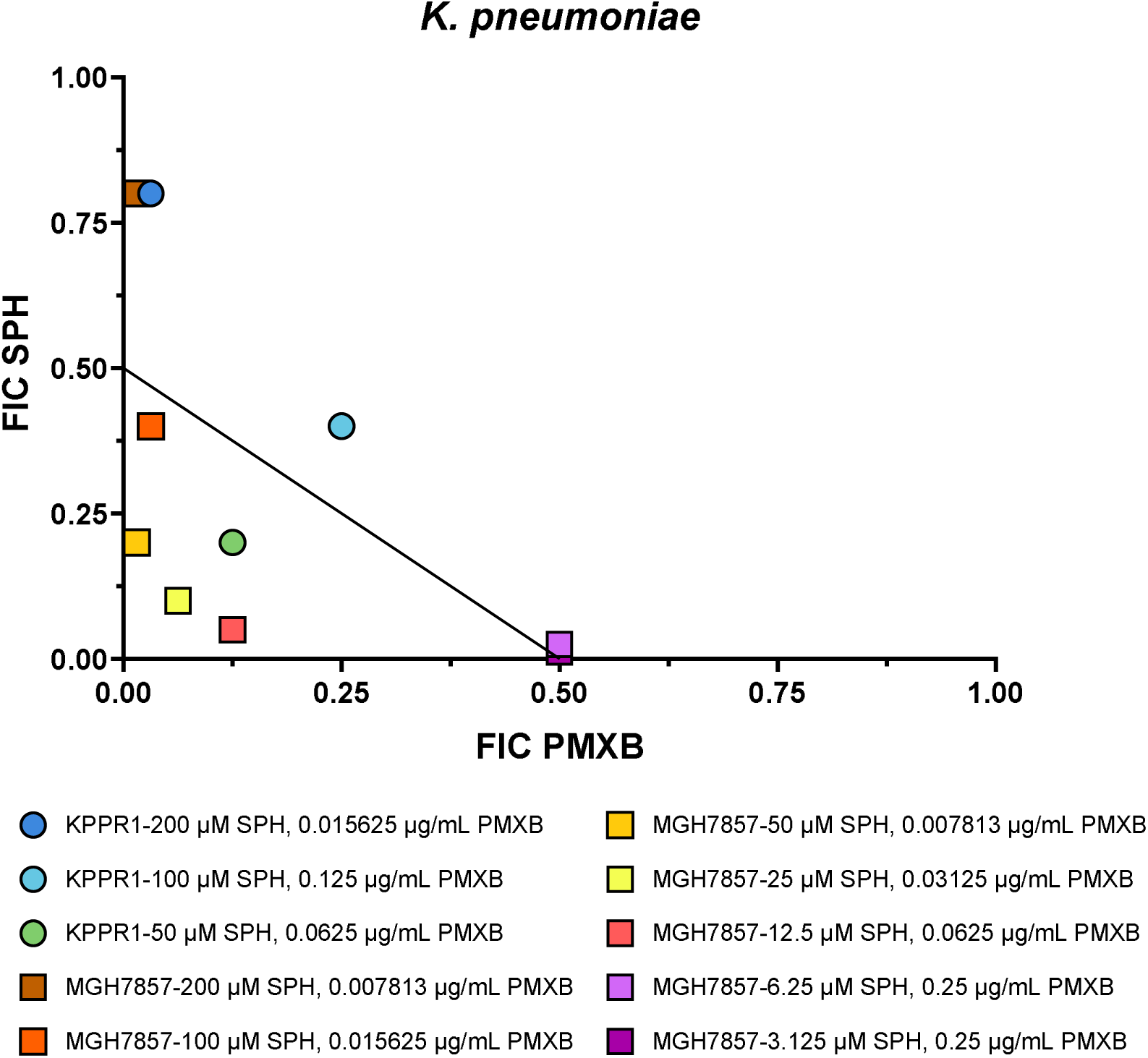
Isobologram of FICI values determined from *K. pneumoniae* checkerboard assays. FIC of sphingosine (FIC SPH) were plotted against the FIC of polymyxin B (FIC PMXB). FICI ≤0.5, >0.5 and ≤ 1, >1 and ≤2, and >2, represent synergistic, additive, unrelated, and antagonistic effects, respectively [46]. Antagonistic effects are not shown for clarity.

Interestingly, the combination of 1 µg/mL polymyxin B and either 6.25 µM or 3.125 µM sphingosine resulted in a slight antagonistic interaction with FICI of 2.025 and 2.0125, respectively (**Figure 7**, **Table 2**).

### Sphingosine and polymyxin combine to enhance killing of *P. aeruginosa* and *K. pneumoniae*

In addition to growth inhibition, we wanted to observe what concentration of polymyxin B and sphingosine resulted in killing of *P. aeruginosa* and *K. pneumoniae*. In our system, sphingosine alone has not displayed bactericidal activity against *P. aeruginosa* and *K.* pneumoniae, while 0.5 µg/mL of polymyxin B is sufficient to kill either bacterium. Against *P. aeruginosa,* the presence of 50 µM sphingosine reduced the minimum bactericidal concentration (MBC) of polymyxin B to 0.031 µg/mL (**Table 3**). We observed that the lowest concentration of polymyxin B needed to kill either strain of *K. pneumoniae* was with the addition of 200 µM sphingosine. With 200 µM sphingosine, the MBC of polymyxin B for MGH78578 and KPPR1 was 0.063 µg/mL and 0.047 µg/mL, respectively (**Table 3**).

**Table 3.** Median MBCs of polymyxin B in combination with sphingosine.

| MBC of Polymyxin B (µg/mL) |  |  |  |
| --- | --- | --- | --- |
| Sphingosine (µM) | *P. aeruginosa* PA14 | *K. pneumoniae* MGH78578 | *K. pneumoniae* KPPR1 |
| 200 | 0.0625 | 0.0625 | 0.046875 |
| 100 | 0.125 | 0.25 | 0.1875 |
| 50 | 0.03125 | 0.15625 | 0.15625 |
| 25 | 0.09375 | 0.25 | 0.5 |
| 12.5 | 0.25 | 0.5 | 0.75 |
| 6.25 | 0.5 | 0.75 | 1 |
| 3.125 | 0.5 | 0.5 | 1 |
| 0 | 0.5 | 0.75 | 0.5 |

## Discussion

The increasing prevalence of multidrug-resistant Gram-negative bacteria is a threat to global health and requires the development of new agents to treat infections caused by these organisms [1, 3, 18]. Although novel antimicrobial therapies are needed, drug development requires an immense amount of capital and time which delays their introduction into the clinic. Alongside the slow progression of drug discovery and development, a reassessment of “retired” antibiotics and the inquiry into effective combinations with potential adjuvants are needed to prolong and enhance the utility of current antibiotics [20].

The purpose of this study was to investigate whether the antibacterial properties of sphingosine could enhance the activity of currently prescribed antibiotics against *P. aeruginosa*. Our initial screen of the effect of sphingosine versus a panel of antibiotics revealed that sphingosine increased the antibacterial activity of polymyxin B and colistin. We observed polymyxin B and colistin synergize with sphingosine at multiple concentrations. The greatest degree of synergy with polymyxin B or colistin occurred with the addition of 50 µM sphingosine, resulting in a FICI of 0.263 and 0.35, respectively. These results were reflected in the enhanced bactericidal effects of polymyxin B when combined with sphingosine, best illustrated by the addition of 50 µM sphingosine, which reduced polymyxin B MBC from 0.5 µg/mL to 0.063 µg/mL. The other sphingoid bases, sphinganine and phytosphingosine, also enhanced polymyxin B activity. The potent antimicrobial activity of phytosphingosine was surprising as we previously reported that phytosphingosine does not affect the growth of *P. aeruginosa* unless the *sphBCD* operon was deleted [38]. Compared to wildtype *P. aeruginosa, sphBCD* deletion mutants display reduced growth in sphingosine, sphinganine, and phytosphingosine with IC_50_ values of 84 µM, 63 µM, and 40 µM, respectively. This discrepancy is likely due to the differences in initial concentration of bacteria. In this study we started *P. aeruginosa* at an OD_600_ of 0.001 (corresponding to 1x10^6^ colony forming units (CFU)/mL) [47], whereas previously we began bacterial cultures at an OD_600_ of 0.05 (5x10^7^ /mL) [38]. Altered antimicrobial activity of sphingosine and phytpsphingosine due to different concentrations of bacteria and sphingoid base have been reported previously [27, 28, 32]. In addition to *P. aeruginosa*, sphingosine enhanced the bacteriostatic and bactericidal activities of polymyxin B against two strains of *K. pneumoniae*, MGH78578 and KPPR1. The enhanced bacteriostatic effects were reflected in bacterial killing, where the addition of 200 µM sphingosine drastically reduced the concentration of polymyxin B needed to kill either strain of *K. pneumoniae*. Interestingly, antagonism between sphingosine and polymyxin B was observed when KPPR1 was exposed to 3.125 µM or 6.25 µM sphingosine, and these results were observed in all checkerboard assays and MBC assays.

To our knowledge, this is the first report of antibiotic synergy between polymyxins and sphingosine affecting the growth and survival of *P. aeruginosa*. Polymyxins, which have been clinically available for about 60 years, have been the focus of numerous studies investigating their potential for combinatorial antibiotic therapies [48–55]. Additionally, the antimicrobial activity of sphingosines has been reported on over the last three decades [9, 26–38, 56–59]. Similar studies have been conducted using a sphingosine-1-phosphate analog, fingolimod (FTY720), and demonstrated that fingolimod has antibacterial activity against Gram-positive bacteria [60]. Recently, a study conducted by Geng *et al*. [55] exhibited antibiotic synergy between fingolimod and colistin against *K. pneumoniae* which inhibited growth of *K. pneumoniae in vitro* and reduced mouse mortality in a murine pneumonia model. The authors hypothesized that the enhanced activity was due to the disruption of bacterial membranes by the combination and inhibition of membrane repair machinery by fingolimod, resulting in a loss of membrane integrity and the leakage of ATP [55]. A similar mechanism has been proposed for the antibacterial activity of sphingosines, where sphingosine induces continuous degradation of cardiolipin, resulting in membrane destabilization while exacerbating repair mechanisms, ultimately leading to death [31, 32]. It is generally understood that polymyxins disrupt outer membranes by electrostatically interacting with lipopolysaccharide (LPS) and displacing the divalent cations (Mg^2+^ and Ca^2+^) for the insertion of its N-terminal fatty acyl group and hydrophobic amino acid side chains into the outer membrane fatty acyl layer [61, 62]. However, there are significant gaps in knowledge of the exact mechanism of action of polymyxins and the role antimicrobial adjuvants play in that mechanism. When paired with sphingosine, the combined interactions of sphingosine with cardiolipin and polymyxins with LPS may exacerbate membrane repair, leading to a rapid loss of membrane integrity. Alternatively, sphingosine may alter outer membrane fluidity to improve polymyxin entry. However, the inability of sphingosine to render *P. aeruginosa* susceptible to vancomycin or substantially reduce the efficacy of aminoglycoside antibiotics (**Table 1, Supplemental Figures 1, 2, and 4**) suggests that sphingosine alone is unable to fully permeabilize the outer or inner membranes, respectively [53, 54]. Therefore, more work is needed to determine the mechanism of sphingosine and polymyxin synergy and convert clinical applications.

Compared to polymyxins alone, the combination of polymyxin and sphingosine permits using significantly lower concentrations of polymyxins to kill Gram-negative bacteria, potentially mitigating adverse effects from polymyxin treatment which has limited their clinical use. In the clinic, polymyxins are delivered intravenously to treat bacteremia or by nebulized inhalation for the treatment of bacterial induced pneumonia, the latter is associated with increased efficacy and reduced systemic adverse effects [63]. Inhalation of nebulized sphingosine has been demonstrated as safe and effective at reducing bacterial loads in porcine pneumonia models *ex vivo* and *in vivo* [56, 58, 59]. Therefore, inhalation of a sphingosine-polymyxin combination may prove to be an effective treatment for bacterial pneumonia. Alternatively, topical application of sphingosine and polymyxins could be used to treat infections in superficial cuts and lacerations, similar to common over-the-counter triple antibiotic ointments, which contain polymyxin B in combination with neomycin and bacitracin. Our data shows that sphingosine and polymyxins synergize to inhibit the growth of *P. aeruginosa* and *K. pneumoniae*; however, a caveat is that our experiments were conducted using a modified MOPS minimal media rather than the CLSI-recommended Mueller-Hinton Broth (MHB). In addition to a defined Mg^2+^ and Ca^2+^ concentration, which would need to be adjusted in MHB [64], minimal media likely better reflects the lower growth rates of bacteria, including *P. aeruginosa*, during infection [65–70]. We have reported previously that the carbon source in minimal media impacts the effect of sphingosine on *P. aeruginosa* growth, where carbon sources enabling faster growth proportionally limit sphingosine’s growth-limiting effects [38]. However, when considering future applications of sphingosine-polymyxin combinations, their effectiveness should be tested in host-mimetic media that simulates the infectious environments of the skin, lung, eye, and ear [71–73].

Overall, this study highlights the potential of sphingosine as an antimicrobial adjuvant for polymyxins against Gram-negative bacteria. The addition of sphingosine dramatically increased the bacteriostatic and bactericidal activities of colistin and polymyxin B against *P. aeruginosa* and *K. pneumoniae*. Although additional studies are needed to determine the combined mechanism of action and the efficacy of the combination in host-mimetic conditions, this pairing of polymyxins and sphingosine represents a prospect for future therapy.

## Materials and Methods

### Strains and growth conditions

*Pseudomonas aeruginosa* (PA14 and PAO1) and *Klebsiella pneumoniae* (MGH78578 and KPPR1) strains were maintained at 37°C on lysogeny-broth (LB) Lennox formulation. In preparation for assays, *P. aeruginosa* was grown overnight in modified MOPS medium [74, 75] supplemented with 20 mM pyruvate and 5 mM glucose, whereas *K. pneumoniae* was grown overnight in modified MOPS medium supplemented with 20 mM lactate and 5 mM glucose.

### Chemicals, antibiotics, and notes on sphingosine stability and solubility

All media and standard chemicals were purchased from ThermoFisher (Waltham, MA) or Sigma-Aldrich (St.Louis, MO). Gentamicin, tobramycin sulfate, aztreonam, ceftazidime, ciprofloxacin, and vancomycin hydrochloride were purchased from ThermoFisher. Polymyxin B sulfate and colistin sulfate were both obtained from MP Biomedicals (Irvine, CA). Antibiotics were stored at -20°C as single-use aliquots to avoid multiple freeze-thaw cycles. The sphingoid bases sphingosine, sphinganine, and phytosphingosine were purchased from Cayman Chemical (Ann Arbor, MI) and dissolved in 95% ethanol to make 50 mM stocks. Sphingoid bases were stored as aliquots at -20°C to avoid multiple freeze-thaw cycles which impact the antibacterial activity of the lipids [9, 38, 76]. Sphingoid bases were administered to the multi-well plates in ethanol and allowed to air dry to remove solvent prior to media addition.

### Combination antimicrobial susceptibility assays

Antimicrobial susceptibility assays to determine MICs were conducted by the broth microdilution method as recommended by the CLSI guidelines [47] with the exception that MOPS media was used instead of Mueller-Hinton Broth (**See Discussion**). Serial twofold dilutions of antibiotics were prepared in modified MOPS medium supplemented with 20 mM of pyruvate or lactate for growth of *P. aeruginosa* and *K. pneumoniae*, respectively, and added to the 96-well plates containing the dried sphingoid bases. Overnight cultures were harvested by centrifugation, washed in MOPS media, and resuspended in MOPS with 20 mM pyruvate for *P. aeruginosa* or 20 mM lactate for *K. pneumoniae* at an OD_600_ of 0.01 or 0.001, respectively.

Adjusted cultures were then added to the 96-well plates containing antibiotic and sphingoid bases at a 1:10 (v:v) ratio for a starting OD_600_ of 0.001 (1x10^6^ CFU/mL) or 0.0001 (1x10^5^ CFU/mL) for *P. aeruginosa* and *K. pneumoniae*, respectively. The difference in initial OD_600_ is to correct for the faster growth rate of *K. pneumoniae* compared to *P. aeruginosa* [77] in these media conditions. To allow for equal gas exchange in each well, 96-well plates were covered with sterile microporous sealing film from USA Scientific (Ocala, FL). Cultures were grown for 18 hours at 37°C with horizontal shaking at 170 rpm. The OD_600_ was measured using a BioTek Synergy H1 plate reader at 0 and 18 hours to assess growth. To assess the change in growth after 18 hours, the OD_600_ at 0 hours was subtracted from the 18-hour reading and the difference was then divided by an untreated control. MICs were determined by the lowest concentration of antibiotic with or without sphingoid base that inhibited ≥90% of growth, relative to an untreated control.

### Checkerboard assays to determine fractional inhibitory concentrations

Similar to MIC assays, for checkerboard assays sphingoid bases were diluted separately to 5 mM, 500 µM, and 250 µM stocks and applied directly to the 96-well plates used for assays to create a range of twofold dilutions from 200 µM to 3.125 µM. After ethanol vehicle had evaporated, twofold serial dilutions of antibiotic in MOPS with 20 mM pyruvate or 20 mM lactate were added to plates containing the dried sphingoid bases. Like the MIC assays, washed and adjusted cultures were added to assay plates at a starting OD_600_ of 0.001 and 0.0001 for *P. aeruginosa* and *K. pneumoniae*, respectively. Assay plates were sealed with breathable microporous film and cultures were grown for 18 hours at 37°C with horizontal shaking at 170 rpm. The OD_600_ was measured using a BioTek Synergy H1 plate reader at 0 and 18 hours to assess growth. To assess the change in growth after 18 hours, the OD_600_ at 0 hours was subtracted from the 18-hour reading and the difference was then divided by an untreated control. MICs were determined by the lowest concentration of antibiotic with or without sphingoid base that inhibited ≥90% of growth, relative to an untreated control. MIC values were used to calculate the fractional inhibitory concentration index (FICI) of the combinations (**Equation 1**). The combined effects of the antimicrobials were characterized as synergistic, additive, unrelated, and antagonistic when the FICI was ≤0.5, 0.5-1.0, 1-2, and >2, respectively [46].

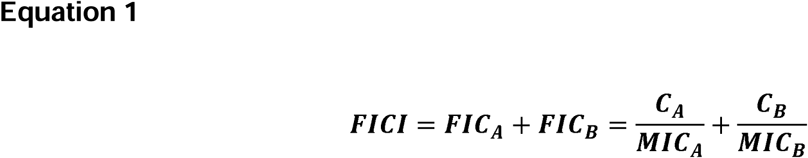

**Equation 1** The fractional inhibitory concentration index FICI, represents the sum of the fractional inhibitory concentrations (*FIC_A_*and *FIC_B_*) of each drug tested, which are determined for each drug by dividing the MIC of each drug when used in combination (*C_A_*and *C_B_*) by the MIC of each drug when used alone (*MIC_A_*and *MIC_B_*).

### Minimal bactericidal concentration (MBC) assays

To observe the bactericidal effects of polymyxins and sphingosine, minimal bactericidal concentration of polymyxin B and sphingosine combinations were determined after checkerboard assays. After 18 hours, 10 µL were removed from each well of the checkerboard and spotted onto *Pseudomonas* Isolation Agar (PIA) or Reasoner’s 2A agar (R2A) for growth of *P. aeruginosa* and *K. pneumoniae*, respectively. MBCs were determined by the lowest concentration of antimicrobials alone or in combination where no colonies were observed. The limit of detection was set at 100 CFU/mL.

### Data acquisition

All data gathered is from three independent experiments, each with technical duplicates.

Results were presented as mean ± standard error of mean. Data collected using a BioTek Synergy H1 were exported to Microsoft Excel, and all normalization and fold change calculations were done in Excel. GraphPad Prism was used for visualization.

## Acknowledgements

We would like to thank Chris Huston for helpful discussions and initial review of this manuscript. This work was supported by funding from the Cystic Fibrosis Foundation to MJW (WARGO24G0).

## Supplemental Figures

**Supplemental Figure 1.**
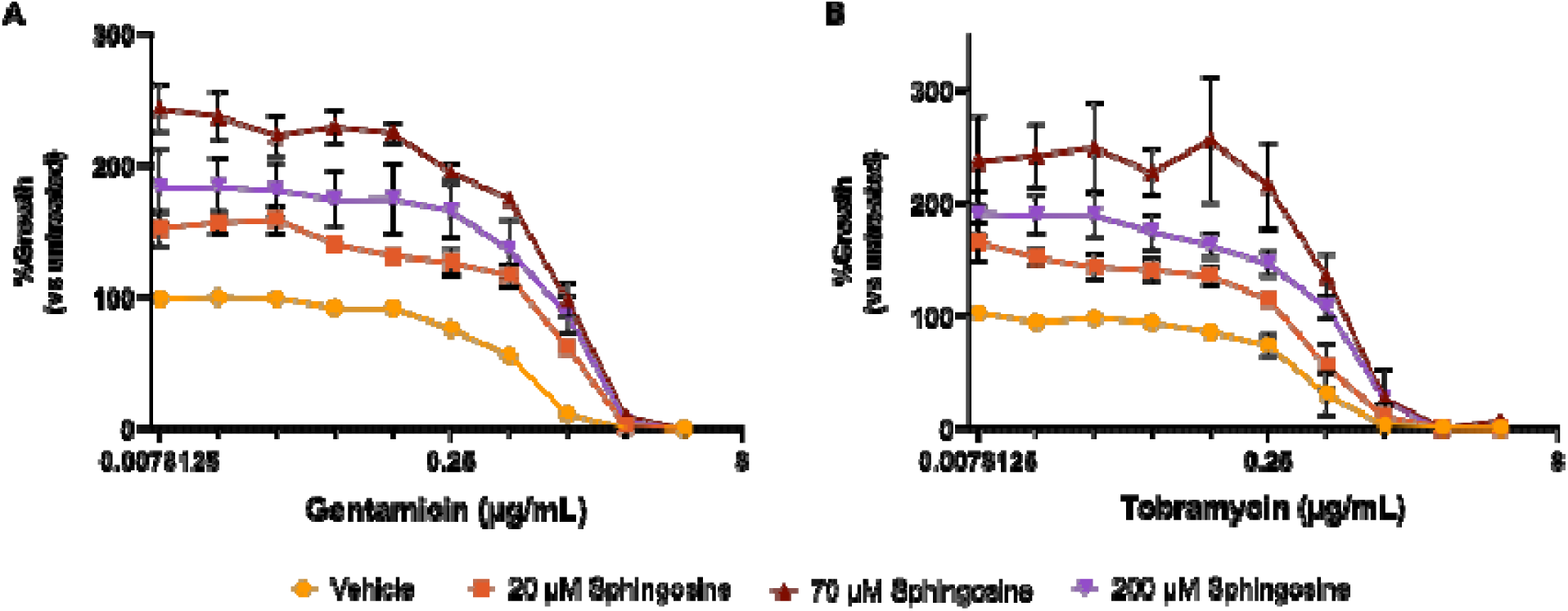
Dose effect curves of PA14 in increasing concentrations of gentamicin **(A)** or tobramycin **(B)** alone or with 20 µM, 70 µM, and 200 µM sphingosine. Data points denote means summarizing three independent experiments and error bars represent the standard error of mean.

**Supplemental Figure 2.**
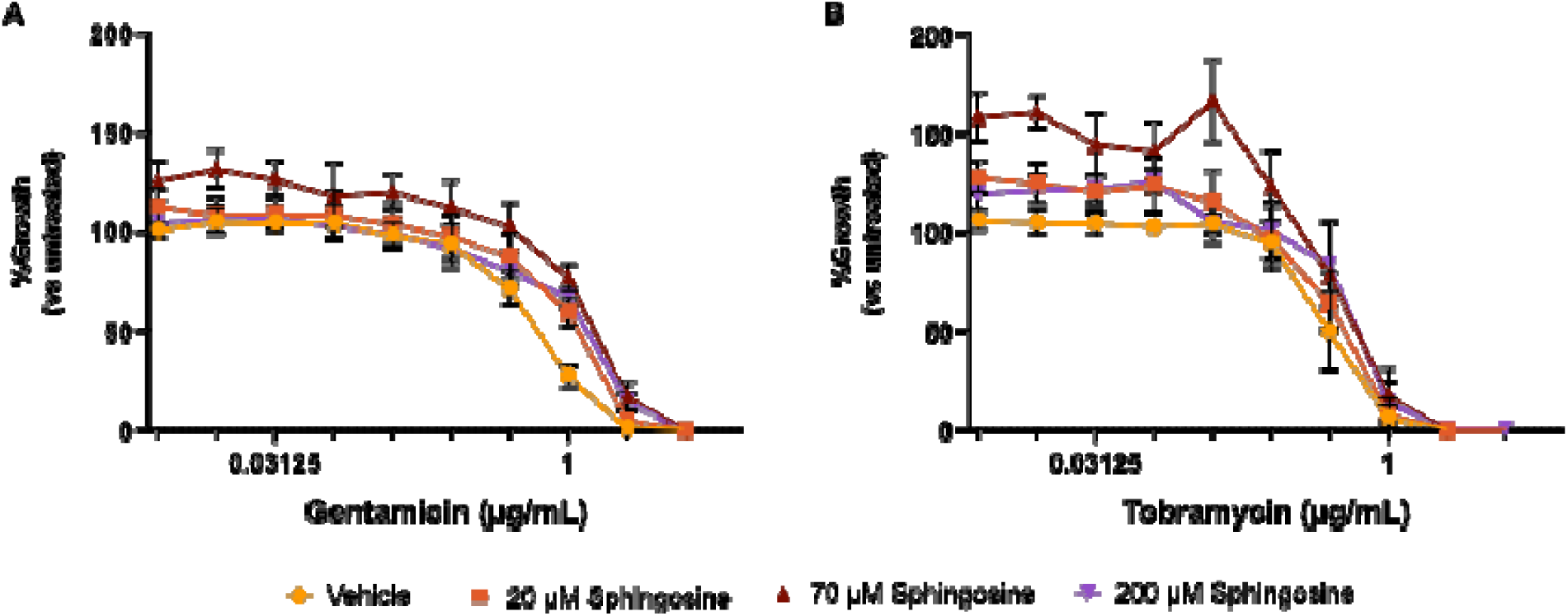
Dose effect curves of PAO1 in increasing concentrations of gentamicin **(A)** or tobramycin **(B)** alone or with 20 µM, 70 µM, and 200 µM sphingosine. Data points denote means summarizing three independent experiments and error bars represent the standard error of mean.

**Supplemental Figure 3.**
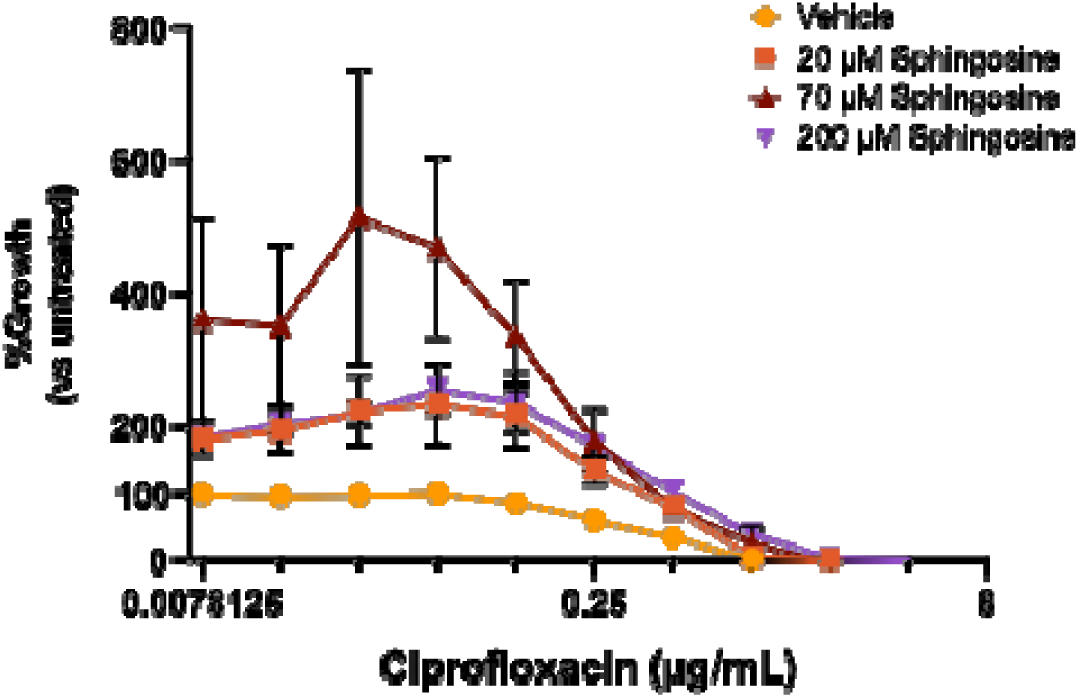
Dose effect curves of PA14 in increasing concentrations of ciprofloxacin alone or with 20 µM, 70 µM, and 200 µM sphingosine. Data points denote means summarizing three independent experiments and error bars represent the standard error of mean.

**Supplemental Figure 4.**
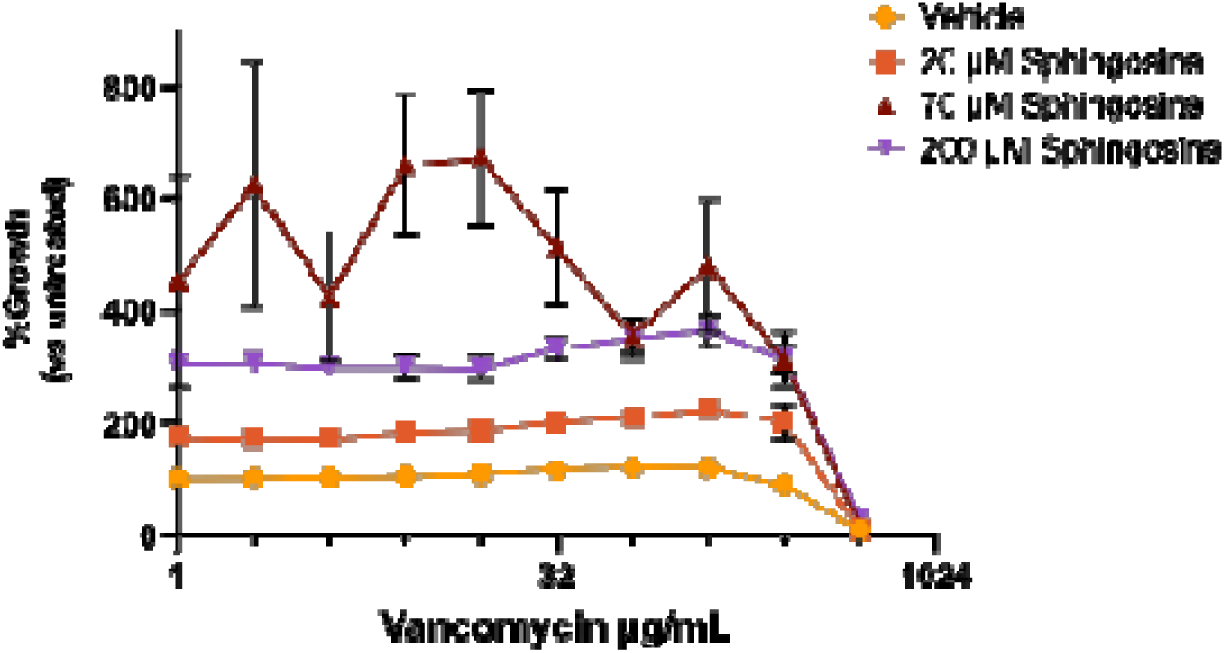
Dose effect curves of PA14 in increasing concentrations of vancomycin alone or with 20 µM, 70 µM, and 200 µM sphingosine. Data points denote means summarizing three independent experiments and error bars represent the standard error of mean.

**Supplemental Figure 5.**
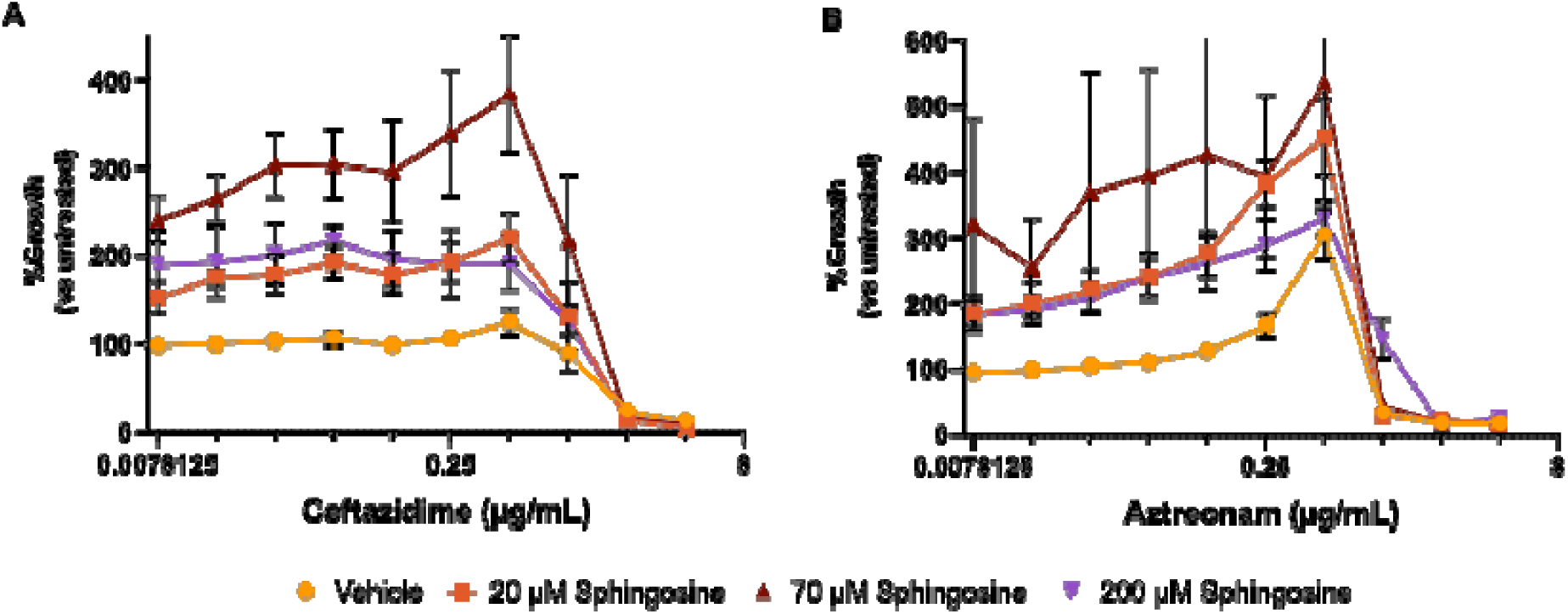
Dose effect curves of PA14 in increasing concentrations of ceftazidime **(A)** or aztreonam **(B)** alone or with 20 µM, 70 µM, and 200 µM sphingosine. Data points denote means summarizing three independent experiments and error bars represent the standard error of mean.

